# Multi-task benchmarking of single-cell multimodal omics integration methods

**DOI:** 10.1101/2024.09.15.613149

**Authors:** Chunlei Liu, Sichang Ding, Hani Jieun Kim, Siqu Long, Di Xiao, Shila Ghazanfar, Pengyi Yang

## Abstract

Single-cell multimodal omics technologies have empowered the profiling of complex biological systems at a resolution and scale that were previously unattainable. These biotechnologies have propelled the fast-paced innovation and development of data integration methods, leading to a critical need for their systematic categorisation, evaluation, and benchmark. Navigating and selecting the most pertinent integration approach poses a significant challenge, contingent upon the tasks relevant to the study goals and the combination of modalities and batches present in the data at hand. Understanding how well each method performs multiple tasks, including dimension reduction, batch correction, cell type classification and clustering, imputation, feature selection, and spatial registration, and at which combinations will help guide this decision. This study develops a much-needed guideline on choosing the most appropriate method for single-cell multimodal omics data analysis through a systematic categorisation and comprehensive benchmarking of current methods.

The Stage 1 protocol for this Registered Report was accepted in principle on 30th July 2024. The protocol, as accepted by the journal, can be found at https://springernature.figshare.com/articles/journal_contribution/Multi-task_benchmarking_of_single-cell_multimodal_omics_integration_methods/26789902.

## Main text

The recent development of single-cell multimodal omics technologies has revolutionised our ability to simultaneously profile multi-layered molecular programs at a global scale in individual cells^1,2^. The availability of single-cell multimodal omics data provides new opportunities to investigate molecular programs that underlie cell identity, cell-fate decisions, and disease mechanisms at a resolution previously inaccessible from studying bulk cell populations^1–3^. A considerable number of technologies and platforms have been developed, which have led to the generation of a variety of combinations of data modalities each capturing a unique molecular feature in the cell. To date, prevalent modalities encompass gene expression (typically denoted as RNA), surface protein abundance (through antibody derived tags [ADT]), and chromatin accessibility (through assay for transposase-accessible chromatin [ATAC]), and can be profiled jointly by popular single-cell multimodal omics technologies such as CITE-seq^4^, SHARE-seq^5^, and TEA-seq^6^ to name a few (Fig. 1a).

**Fig. 1:**
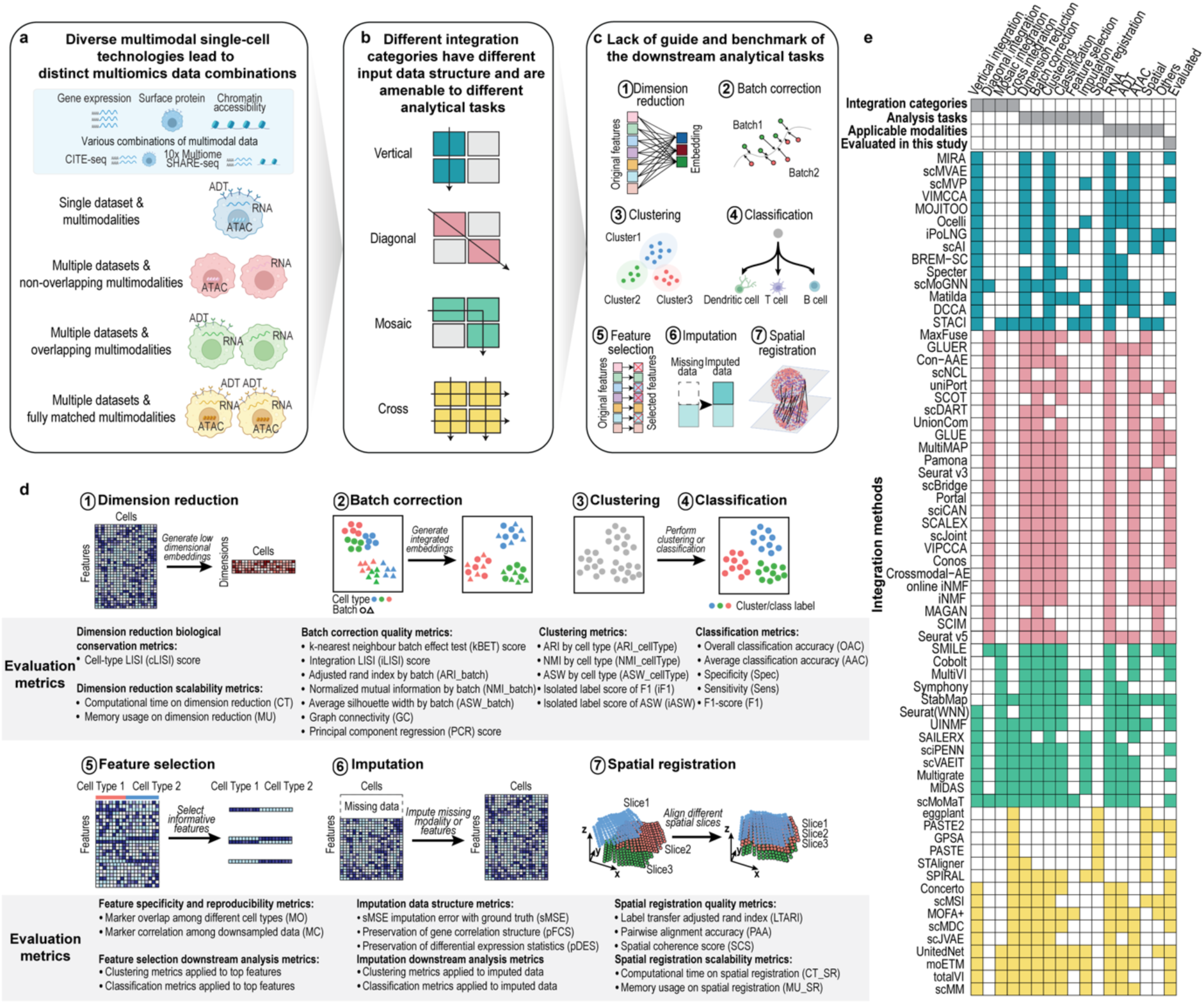
An overview of benchmarking for single-cell multimodal omics integration methodologies. **a**, Diverse technological combinations arising from different modalities. **b**, Four prototypical single-cell multimodal omics data integration categories. **c**, Schematics of common tasks for single-cell multimodal omics analysis. **d**, Schematics for each task and the panel of evaluation metrics tailored for its performance evaluation. **e**, Summary of integration methods and their alignment with specific integration categories, computational tasks, modality capacities, and evaluation status.

Integrating modalities of data generated from single-cell multimodal omics technologies is essential and greatly impacts the utility of such data for downstream biological interpretation^7–9^. A fast-growing number of computational methods have been developed to integrate various combinations of data modalities captured by different single-cell multimodal omics technologies. Nevertheless, there is a lack of study that comprehensively outlines the similarities and differences between current methods including the data integration strategies, the tasks they intend to address, and the data modalities they can integrate. Furthermore, while a few works have been conducted to compare integration methods for cancer studies using both bulk and single-cell omics data^10,11^, there is a lack of systematic evaluation and benchmark of the currently available methods specifically designed for single-cell multimodal omics data integration for their performance on the broad range of tasks and data modalities they are designed to handle. These, together, have created a significant challenge for navigating and selecting the most pertinent integration method for single-cell multimodal omics data analysis.

To address the challenge of selecting the most appropriate methods for single-cell multimodal omics data analysis, here we set out to systematically categorise and comprehensively benchmark the current integration methods on common computational tasks and data modalities. Building on previous works^7,12^ and based on input data structure and modality combination, we define four prototypical single-cell multimodal omics data integration categories, including “vertical”, “diagonal”, “mosaic”, and “cross” integration (Fig. 1b).

Depending on the applications, we further introduce seven common tasks that many methods are designed to address (Fig. 1c), including (1) dimension reduction, (2) batch correction, (3) clustering, (4) classification, (5) feature selection, (6) imputation, and (7) spatial registration. Using panels of evaluation metrics each tailor-made for a specific task (Fig. 1d), we evaluated 40 integration methods (Fig. 1e) across the four data integration categories on 64 real datasets and 22 simulated datasets. In particular, we included 18 vertical integration methods, 14 diagonal integration methods, 12 mosaic integration methods, and 15 cross integration methods. This registered report provides a comprehensive view of methods designed for single-cell multimodal omics data integration, and serves as (i) a much-needed guide for single-cell integration across modalities to the research community and (ii) a foundation to foster future research and application of computational methods for single-cell multimodal omics data analysis.

The study was conducted in accordance with the registered, peer-reviewed protocol at https://springernature.figshare.com/articles/journal_contribution/Multi-task_benchmarking_of_single-cell_multimodal_omics_integration_methods/26789902. Except for pre-registered data, all analysis results reported in the paper were collected after the date of the registered protocol publication.

## Results

### Vertical integration for dimension reduction and clustering

To evaluate the performance of vertical integration methods on dimension reduction and clustering tasks, we systematically benchmarked all applicable methods and datasets of varying modalities. This included evaluations of 14 methods^13–26^ on 13 paired RNA and ADT (RNA+ADT) datasets, 14 methods^17–30^ on 12 paired RNA and ATAC (RNA+ATAC) datasets, and 5 methods^17–21^ on 4 datasets containing all three modalities (RNA+ADT+ATAC).

As an illustrative example, the results from different methods on a representative dataset D7, with paired RNA and ADT data, are visualised in Fig. 2a and quantitatively summarised in Fig. 2b. Among the evaluated methods, Seurat WNN^22^, sciPENN^14^, and Multigrate^19^ demonstrated generally better performance on this dataset, effectively preserving the biological variation of cell types. While the evaluation metrics generally agreed in method assessment, notable differences in ranking were observed. For instance, moETM^25^ was one of the top ranked methods by iF1 and NMI_cellType, yet received comparatively low rankings based on ASW_cellType and iASW. UMAP visualisations and metric quantifications for additional representative datasets, D15 (RNA+ATAC) and D22 (RNA+ADT+ATAC), are shown in Supplementary Fig. 1a,b and Supplementary Fig. 1c,d, respectively. We note that Seurat WNN, MIRA, and scMoMaT generate graph-based outputs rather than embeddings, making ASW_cellType and iASW metrics inapplicable for these methods.

**Fig. 2:**
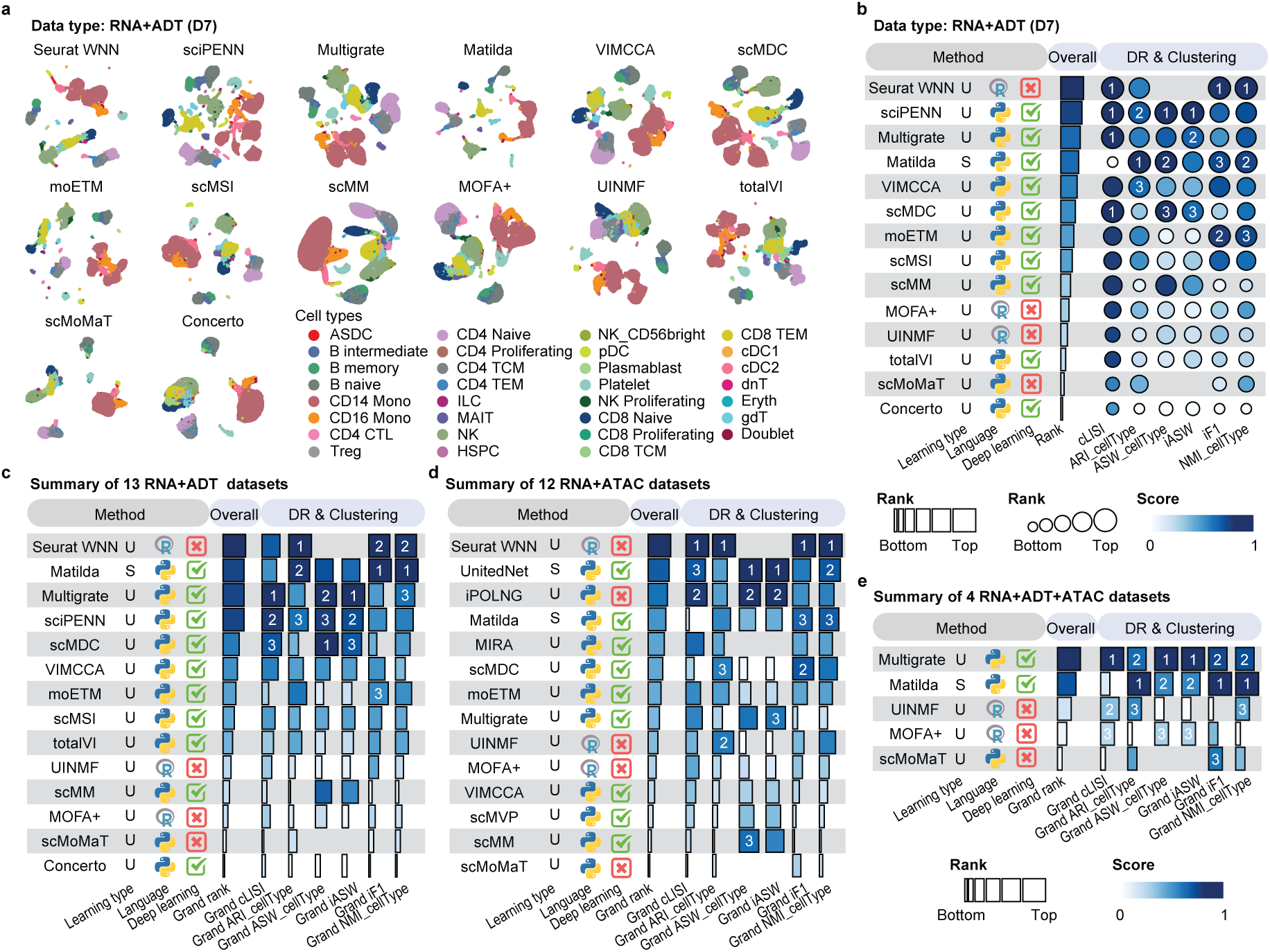
Benchmark results of vertical integration methods for dimension reduction and clustering. **a**, UMAP visualisation of the vertical integration methods applied to a representative RNA+ADT dataset (D7). **b**, Method performance on the dataset D7. Dimension reduction (DR) and clustering metrics are used to evaluate method performance. Overall rank scores (horizontal bars) are computed as the min-max scaled mean ranks across evaluation metrics. Individual metric scores (bubbles) are computed as the min-max scaled raw values. Performance summary of vertical integration methods from **c**, All RNA+ADT datasets; **d**, All RNA+ATAC datasets; and **e**, All RNA+ADT+ATAC datasets. Overall grand rank scores are computed as the min-max scaled mean rank of the grand ranks across each metric. Grand ranks of individual metrics are computed as the min-max scaled mean ranks across all applicable datasets. For all summary panels, the learning types are categorised into three groups including supervised (S), unsupervised (U), and semi-supervised (Semi). Metrics that do not apply to a given method were not assigned a value.

To summarise the method performance across all datasets for the dimension reduction and clustering tasks, we first calculated the overall rank score for each dataset by summarising method performance across evaluation metrics (Supplementary Fig. 1e-g). Boxplots of ranks for each method visualise the variation in method performance across individual datasets (Supplementary Fig. 1h–j). Methods were then ranked based on their overall grand rank scores for 13 bimodal datasets of RNA+ADT (Fig. 2c), 12 bimodal datasets of RNA+ATAC (Fig. 2d), and 4 trimodal datasets of RNA+ADT+ATAC (Fig. 2e). These analyses reveal that dataset complexity may affect the performance of integration methods. Simulated datasets, which may lack latent data structure observed in real data, can be easier to integrate and some methods (e.g. scMM^23^) that do not perform well on real datasets tend to perform better on simulated datasets (Supplementary Fig. 1e-f). While Seurat WNN, Multigrate, Matilda^17^, and UnitedNet^29^ in general performed well across diverse datasets for particular data modality combinations (Supplementary Fig. 1e-f), method performance is dataset- and, more notably, data modality-dependent.

### Benchmark vertical integration for feature selection

Feature selection is typically used to identify molecular markers associated with specific cell types^31^. Among the vertical integration methods, only Matilda^17^, scMoMaT^21^, and MOFA+^18^ support feature selection of molecular markers from single-cell multimodal omics data.

Notably, Matilda and scMoMaT are capable of identifying distinct markers for each cell type in a dataset, whereas MOFA+ selects a single cell-type-invariant set of markers for all cell types. Extended Data Fig. 1a visualises the top cell-type-specific markers selected from the RNA and ADT modalities of dataset D8 (RNA+ADT) for three example cell types (i.e. “CD14 Mono”, “NK”, and “Plasmablast”) by Matilda and scMoMaT. MOFA+ is excluded from this visualisation as its marker selection does not associate with a particular cell type.

For both Matilda and scMoMaT, the selected markers show higher gene expression (RNA) and protein abundance (ADT) in their respective cell types compared to other cell types.

Moreover, for RNA of NK cells, ADT of CD14 monocytes, and ADT of Plasmablast cells, the two methods identified the same top markers. The quantification of method performance on feature selection for each data modality, using the union of top-5 markers selected from all cell types, in D8 is presented in Extended Data Fig. 1b. To ensure comparability, the number of markers selected by MOFA+ was matched with the other two methods. Results from D8 suggest that the assessments of selected markers can differ across clustering and classification metrics. Visualisation and quantification of feature selection results from additional representative datasets with varying combinations of data modalities are shown in Extended Data Fig. 2a-d, which demonstrates more consistent evaluation results across clustering and classification metrics.

To summarise the method performance across all datasets for feature selection task, we first calculated the overall rank score of each method in each dataset for the three evaluation categories of clustering, classification, and reproducibility for datasets with RNA+ADT modalities (Supplementary Fig. 2a), RNA+ATAC modalities (Supplementary Fig. 2b) and RNA+ADT+ATAC modalities (Supplementary Fig. 2c). Variations in method performance across individual datasets are visualised in Extended Data Fig. 2e,f and Supplementary Fig. 2d. Methods were then ranked based on their overall grand rank scores for each combination of data modalities (Extended Data Fig. 1c-e). These results reveal that MOFA+, while unable to select cell-type-specific markers, generated more reproducible feature selection results across different data modalities. Note that marker correlation (MC) is not influenced by which top features are chosen. Instead, it measures the correlation across all features, leading to the same values even when the selected markers vary within the same method. However, features selected by scMoMaT and Matilda generally led to better clustering and classification of cell types than those by MOFA+. It is also interesting to note that markers selected from ATAC modality (i.e. peaks) typically underperform in clustering and classification evaluations compared to those selected from RNA and ADT modalities (Supplementary Fig. 2b,c). Furthermore, increasing the number of top markers did not always improve downstream evaluation results. Importantly, method performance varies depending on the datasets and evaluation metrics, underscoring the necessity of using multiple evaluation metrics for robust method assessment.

### Benchmark diagonal integration methods

We systematically benchmarked 14 diagonal integration methods^32–45^ on dimension reduction, clustering, batch correction, and classification tasks, including evaluations on 12 datasets with a single batch of RNA data and a single batch of ATAC data, and 5 methods^40–44^ on 6 datasets comprising multiple batches of RNA data and multiple batches of ATAC data. As an example, the evaluation results of different methods on a representative dataset D27, which consists of one RNA batch and one ATAC batch, are visualised in Fig. 3a and quantified using different evaluation metrics in Fig. 3b. Among the methods, scBridge^32^, GLUE^44^, Seurat v5^38^, uniPort^45^, and scJoint^43^ achieved the best overall performance on this dataset, effectively removing batch effects while preserving the biological variation of cell types. Of note, GLUE and Seurat v5 integrate RNA and ATAC data using peak features, whereas the other methods require transforming ATAC peaks into gene activity scores. While the evaluation metrics largely agreed in method assessment, some differences in ranking were observed on this dataset. For instance, scBridge was top-ranked in dimension reduction and clustering whereas the other high-performing method GLUE was top-ranked in batch correction. Moreover, rankings of methods in the classification category further deviate from those in dimension reduction and clustering and batch correction, and could be confounded by the use of the MLP classifier implemented in this work and, if available, the original classifiers implemented in the methods. Results from a representative dataset D37 with multiple batches of RNA and ATAC data are shown in Supplementary Fig. 3a,b. Fewer methods can handle such data, and among those applicable, GLUE and scJoint demonstrated the best performance on this dataset.

**Fig. 3:**
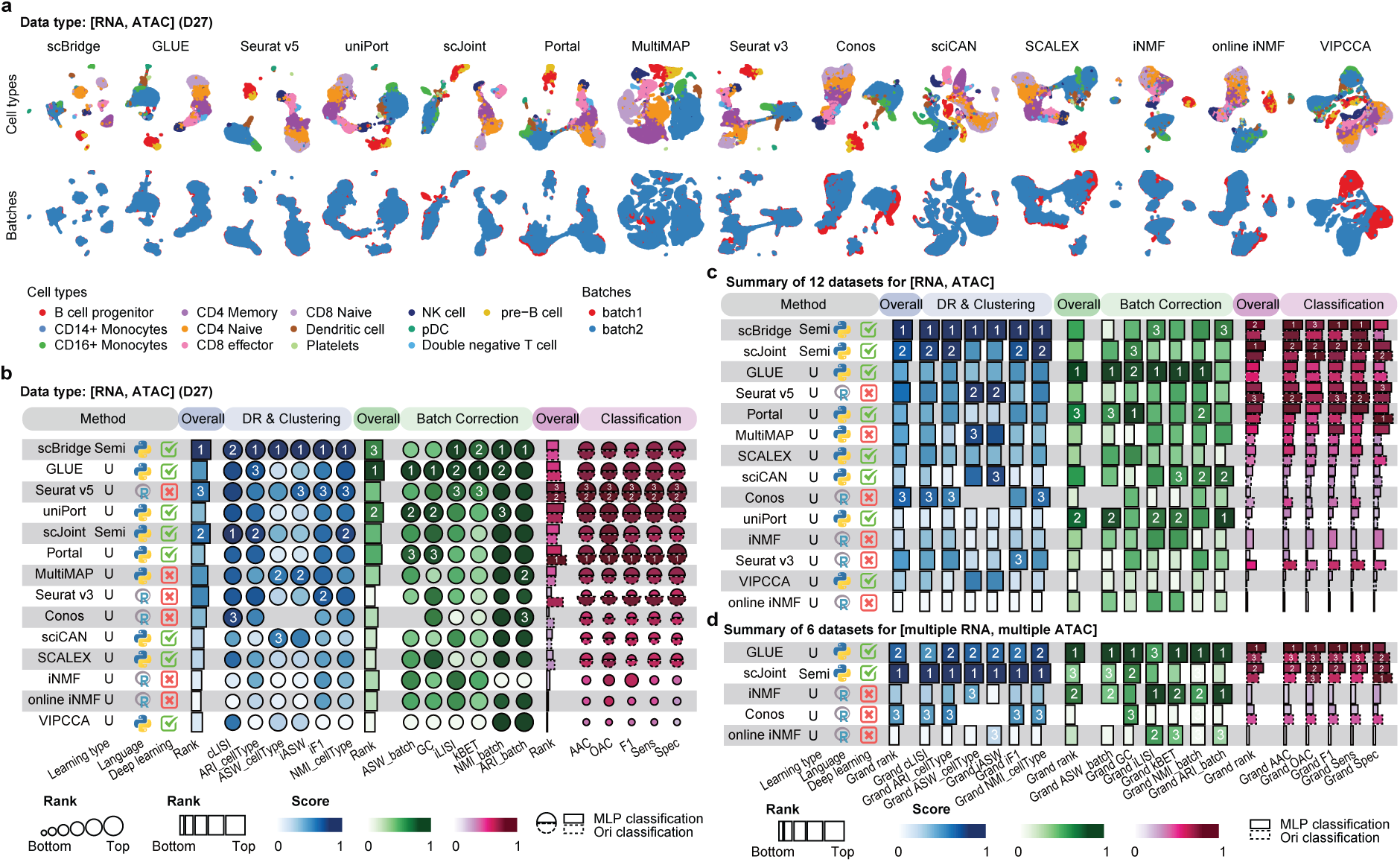
Benchmark results of diagonal integration methods. **a**, Visualisation of diagonal integration methods applied to a representative dataset of D27 with a single batch of RNA data and a single batch of ATAC data. **b**, Method performance on the dataset D27. Metrics are categorised into dimension reduction (DR) and clustering (blue tab), batch correction (green tab), and classification (pink tab). For classification, the solid line represents the MLP classifier implemented in this work, while the dashed line represents the classifier proposed in the original papers. Performance summary of diagonal integration from **c,** All datasets with a single batch of RNA data and a single batch of ATAC data, and **d,** All datasets with multiple batches of RNA data and multiple batches of ATAC data.

To summarise the method performance across all datasets for the dimension reduction, clustering, batch correction, and classification tasks, we first calculated the overall rank score for each dataset (Supplementary Fig. 3c,d) and visualised the performance variability across datasets (Supplementary Fig. 3e,f), and then ranked methods based on their overall grand rank scores for the datasets with single RNA and ATAC batches (Fig. 3c), and for datasets with multiple RNA and ATAC batches (Fig. 3d). These results reveal that methods such as scBridge, and scJoint consistently demonstrated top performance in dimension reduction and clustering tasks across datasets. However, their effectiveness in batch correction was moderate, suggesting limitations in their ability to fully harmonise batches. In contrast, methods like GLUE, uniPort, and Portal^33^, which excel at batch correction, did not demonstrate similarly strong performance in dimension reduction and clustering. This observation indicates a potential trade-off between preserving biological signals, captured by dimension reduction and clustering results, and effective batch correction. Additionally, except for Seurat v3^36^, which performed better with its specifically designed classifier, most integration methods showed similar overall classification rank scores regardless of the classifier used, indicating that the performance depends more on the quality of the integration method itself rather than the choice of classifier. This highlights that, while classifier choice may influence results for some methods, the effectiveness of the integration method is the primary determinant of classification performance.

Taken together, in datasets with single RNA and ATAC batches, scBridge was the top performer across all metrics for dimension reduction and clustering, while GLUE excelled in batch correction metrics. In contrast, for datasets with multiple RNA and ATAC batches, scJoint ranked highest across all dimension reduction and clustering metrics, with GLUE ranking second and demonstrating strong performance on batch correction metrics, highlighting its effectiveness in addressing batch effects.

### Benchmark mosaic integration methods

We next benchmarked seven mosaic integration methods (i.e. MultiVI^46^, Cobolt^47^, SMILE^48^, scMoMaT^21^, Multigrate^19^, StabMap^49^, UINMF^20^) for dimension reduction, clustering, batch correction, and classification tasks, on 17 datasets spanning four data types. As an illustrative example, Fig. 4a visualises UMAP projections of results generated by different methods on dataset D45, which comprises three batches, including an RNA batch, an ATAC batch, and an RNA+ATAC batch serving as a bridge between data modalities. Method performance on this dataset was quantified in Fig. 4b. Among the methods, MultiVI, Cobolt, and SMILE are specifically designed for this type of data. In contrast, scMoMaT, Multigrate, and StabMap are more flexible, as they only require the presence of bridge features between different batches for any data modality combinations. Results from additional datasets with other data modality combinations are presented in Supplementary Fig. 4a-f. In particular, only scMoMaT, Multigrate, and StabMap were applicable to D40, a dataset with three batches (RNA, ADT, RNA+ADT), and D49 (RNA+ADT, RNA+ATAC, ADT). Beside these three methods, UINMF was also applicable to D46 (RNA+ADT, RNA+ATAC, RNA). UINMF is unique in that it requires common features across all batches and therefore is limited in its applicability.

**Fig. 4:**
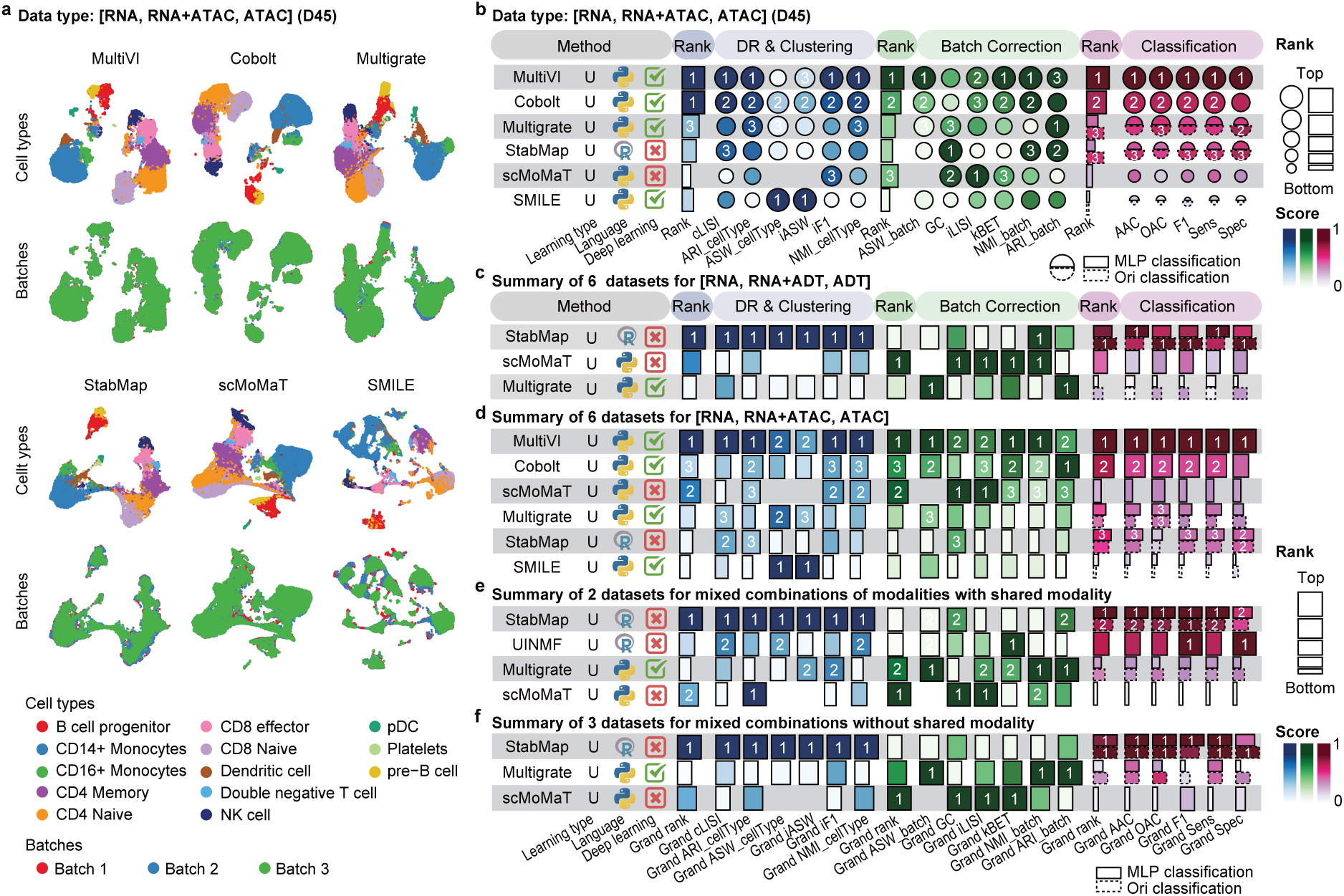
Benchmark results of mosaic integration methods. **a**, UMAP visualisation of mosaic integration methods applied to a representative dataset D45 with data batches of RNA, paired RNA+ATAC, and ATAC. **b**, Method performance on the dataset D45. Metrics are categorised into dimension reduction (DR) and clustering (blue tab), batch correction (green tab), and classification (pink tab). For classification, the solid line represents the MLP classifier implemented in this work, while the dashed line represents the classifier proposed in the original papers. Performance summary of mosaic integration from **c**, All datasets with data batches of RNA, paired RNA+ADT, and ADT; **d**, All datasets with data batches of RNA, paired RNA+ATAC, and ATAC; **e**, All datasets with mixed batch combinations with shared modality across batches; and **f**, All datasets with mixed batch combinations but without shared modality across batches.

To evaluate the overall performance of mosaic integration methods, we first ranked the methods based on their overall rank scores in each dataset and within each task category. In Supplementary Fig. 4g, datasets with (RNA, RNA+ADT, ADT) and those with mixed combinations with or without a shared modality are combined into a single panel, while datasets with (RNA, RNA+ATAC, ATAC) are shown separately in Supplementary Fig. 4h. The overall grand rank scores are presented in Fig. 4c-f, where Fig. 4c summarises results for datasets with (RNA, RNA+ADT, ADT), Fig. 4d presents results for datasets with (RNA, RNA+ATAC, ATAC), and Fig. 4e,f summarise results for datasets with mixed combinations of modalities and with or without shared features. The method performance variability across individual datasets is visualised in Supplementary Fig. 4i,j. These results reveal that StabMap performed well especially in dimension reduction, clustering, and classification tasks across most datasets. scMoMaT and Multigrate show strong performance in batch correction across datasets and data types. When analysing datasets with (RNA, RNA+ATAC, ATAC), MultiVI and Cobolt consistently achieve strong performance across tasks and datasets, likely because they are specifically designed for such data type.

Overall, these analyses provide guidance on method selection for specific tasks and input data types, but also highlight that there are relatively few mosaic integration methods available compared with other integration categories, as many of them impose strict restrictions, limiting their flexibility. This lack of flexibility can make these methods less suitable for datasets with diverse modalities or limited shared features, highlighting the need for more flexible approaches in mosaic integration.

### Benchmark methods on data imputation for mosaic data

While the mosaic integration methods benchmarked in the previous section can be directly applied to mosaic data, several additional methods can also be applied to mosaic data by first imputing missing data modalities and subsequently performing integrative analysis. These include sciPENN^14^, moETM^25^, scMM^23^, totalVI^13^, and UnitedNet^29^. Notably, MultiVI^46^ and StabMap^49^ also offer options for data imputation although they are applicable to mosaic data integration without the imputation step. Among these methods, totalVI and sciPENN can only impute the ADT modality, whereas the other methods can impute both RNA and another modality (ADT or ATAC).

Extended Data Fig. 3a-c presents imputation results, where we imputed the missing ADT modality in a representative CITE-seq dataset D53. UnitedNet and MultiVI were excluded from this analysis as they do not support imputation in this scenario. On this dataset, we found that while data imputed by scMM showed good structural similarity to the ground truth, this did not translate to better clustering and classification results. Conversely, data imputed by sciPENN showed good performance when assessed by clustering and classification metrics but have the lowest structural similarity to the ground truth compared to those imputed by other methods. Similar results were found from additional imputation analyses. These include imputing the RNA modality for the CITE-seq dataset D53 (Supplementary Fig. 5a-c) and imputing ATAC (Supplementary Fig. 5d-f) and RNA (Supplementary Fig. 5g-i) modalities for the 10x multiome dataset D56. These findings highlight that strong preservation of structural similarity during imputation does not necessarily lead to improved performance on downstream analytical tasks, emphasising the importance of evaluating imputation methods across multiple complementary metrics.

To summarise the method performance across all datasets for the imputation task, we first calculated the overall rank score of each method in each dataset for the evaluation categories of clustering, classification, and structural similarity (Supplementary Fig. 5j-m) and visualised their variability in Supplementary Fig. 5n-q. Methods were then ranked based on their overall grand rank scores for each imputation scenario, including imputing ADT (Extended Data Fig. 3d) and RNA (Extended Data Fig. 3e) for CITE-seq datasets, and imputing ATAC (Extended Data Fig. 3f) and RNA (Extended Data Fig. 3g) for multiome datasets. These results revealed that totalVI and StabMap imputed ADT data performed well in clustering and classification evaluations, and demonstrated strong structural similarity to those in ground truth data. Although data imputed by sciPENN achieved the best performance in clustering and classification, their structural similarity did not rank as high as the others. For imputing ATAC or RNA data from 10x multiome, MultiVI demonstrated good performance under clustering and classification metrics, while StabMap and scMM performed well under structural similarity metrics. These analyses highlight that imputation performance alone does not determine the overall effectiveness of a method in mosaic data analysis, and a comprehensive evaluation across multiple criteria is essential.

### Benchmark cross integration methods

We next benchmarked methods for dimension reduction, clustering, batch correction, and classification tasks on cross integration where all data modalities are present in each of all batches in a dataset. This included seven multi-batch bimodal RNA+ADT datasets, four multi-batch bimodal RNA+ATAC datasets, three multi-batch bimodal ADT+ATAC datasets, and three multi-batch trimodal RNA+ADT+ATAC datasets. Results from different methods on a representative dataset D52 are visualised in Fig 5a and quantified by evaluation metrics in Fig. 5b. These results highlight the agreements and differences in performance assessment depending on the evaluation metrics. In particular, performance rankings from dimension reduction, clustering, and classification metrics tend to have higher concordance on this dataset, whereas those from batch correction metrics tend to lead to different rankings.

**Fig. 5:**
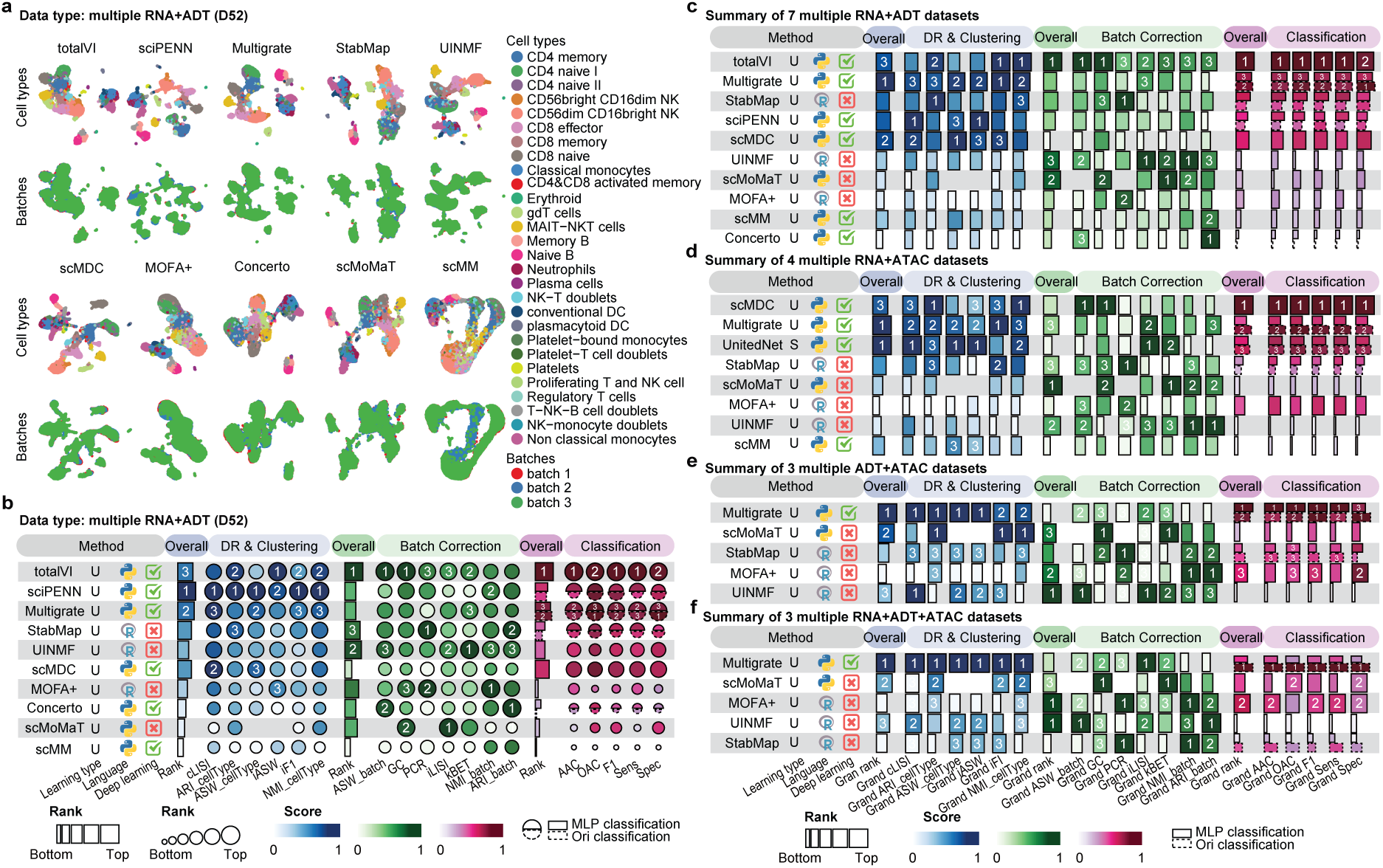
Benchmark results of cross integration methods. **a**, UMAP visualisation of cross integration methods applied to a representative dataset of D52. **b**, Method performance on the dataset D52. Metrics are categorised into dimension reduction (DR) and clustering (blue tab), batch correction (green tab), and classification (pink tab). For classification, the solid line represents the MLP classifier implemented in this study, while the dashed line represents the classifier proposed in the original papers. Performance summary of cross integration methods applied to **c**, All datasets with paired RNA+ADT data across multiple batches; **d**, All datasets with paired RNA+ATAC data across multiple batches; **e**, All datasets with paired ADT+ATAC data across multiple batches; and **f**, All datasets with paired RNA+ADT+ATAC data across multiple batches.

Similar results were found in additional datasets including D56 (multi-batch RNA+ATAC), D58 (multi-batch ADT+ATAC), and D59 (multi-batch RNA+ADT+ATAC) in Supplementary Fig. 6a-f.

We summarised the method performance in each dataset by their overall rank scores (Supplementary Fig. 6g-l) and then ranked methods by their grand overall rank scores across dataset categories, including those from RNA+ADT datasets (Fig. 5c), RNA+ATAC datasets (Fig. 5d), ADT+ATAC datasets (Fig. 5e), and RNA+ADT+ATAC datasets (Fig. 5f). These analyses reaffirmed the above-observed discordance between batch correction metrics and other evaluation metrics, highlighting the challenge of achieving balanced performance across tasks. For example, scMDC performed well in dimension reduction, clustering, and classification, but was less competitive in batch effect removal. Conversely, scMoMaT showed strong performance in batch effect removal but in many cases was less competitive in other tasks (Supplementary Fig. 6g-i). Overall, for bimodal RNA+ADT datasets, totalVI, a method specifically designed for such type of data, performed well across most evaluation metrics (Fig. 5c). For datasets with other combinations of data modalities, Multigrate appears to be the best option given its ability to handle more complex data modality combinations and consistently high performance across various evaluation metrics (Fig. 5d-f).

### Benchmark spatial registration methods

Spatial registration aligns spatial transcriptomics data from different slices into a common spatial framework. Spatial coordinates can be viewed as an additional data modality and its integration conceptually aligns with other integration tasks. To compare method performance on spatial registration for spatial transcriptomics data integration, we benchmarked five methods (i.e. PASTE centre alignment^50^, PASTE pairwise alignment^50^, SPIRAL^51^, GPSA^52^, PASTE2^53^) on 12 datasets generated from various patients/donors and spatial techniques (i.e. Visium, MERFISH, Stereo-seq, Xenium). An example data D61 generated using Visium technology is shown in Extended Data Fig. 4a, where each column visualises the four slices of the spatial transcriptomics profiles of the human cortex aligned by a method. Extended Data Fig. 4b quantifies the alignment results from different methods using three evaluation metrics and then ranks the methods based on the overall performance across these evaluation metrics. Among the evaluated methods, PASTE with centre and pairwise alignments and PASTE2 rely solely on linear transformation, and therefore, preserve the relative spatial relationship within each slice. As a result, their spatial coherence scores (SCS), which quantifies the spatial coherence of tissue layers across different slices, are identical. In contrast, SPIRAL and GPSA apply non-linear transformation to the original slices, and this may explain their relatively low performance when assessed by SCS. Alignment results and their quantification from another representative dataset D64 generated by the MERFISH technology are shown in Supplementary Fig. 7a,b.

The overall rankings for each individual dataset are summarised in Supplementary Fig. 7c, while the grand overall ranking across all datasets is presented in Extended Data Fig. 4c.

Performance variability across individual datasets are summarised in Supplementary Fig. 7d. These results indicate that PASTE with centre alignment performed consistently well across all datasets. PASTE2 also showed strong and stable performance, ranking competitively in most cases. However, its added flexibility in partial alignment across more diverse spatial transcriptomics datasets may offer limited benefit compared to PASTE in the alignment of adjacent and serial tissue sections. In comparison, the performance of PASTE with pairwise alignment, GPSA, and SPIRAL was more variable and appeared to depend on the specific dataset (Supplementary Fig. 7c). It is important to note that the SCS metric yields identical scores for methods that apply linear transformations of each slice, and SPIRAL, which employs a non-linear transformation that regenerates coordinates for query slices, was not favourably evaluated under SCS (Extended Data Fig. 4c). However, SPIRAL demonstrated competitive performance under other metrics such as label transfer adjusted Rand index (LTARI) and pairwise alignment accuracy (PAA). Together, these findings highlight the importance of evaluating spatial registration methods using a broad range of datasets and diverse metrics, given that method performance can depend heavily on both the nature of the data and the chosen assessment criteria.

### Computation time and peak memory usage

As the number of cells in a dataset increases, computational speed becomes a critical factor in integration. To evaluate computational efficiency, we monitored the time and peak memory usage of each method on a selection of large datasets with different data modality combinations for each integration category (Supplementary Fig. 8). Among the methods used for vertical integration, we found that UINMF was one of the most efficient methods both in terms of computation time (Supplementary Fig. 8a) and peak memory usage (Supplementary Fig. 8b) across different data modality combinations and scales well with increasing cell numbers. In contrast, some methods did not scale well with increasing numbers of cells (e.g. scMSI, MIRA) and others consumed a large amount of memory (e.g. scMM, MOFA+, UnitedNet). For diagonal integration, while most methods performed very fast with minimal memory consumption (Supplementary Fig. 8c,d), Seurat appeared to consume significantly more time and memory. For mosaic integration, MultiVI showed a significant increase in computation time usage and scMoMaT showed a significant increase in memory usage (Supplementary Fig. 8e,f). Besides these two methods, Multigrate also required relatively longer computational time and used more memory in most applications. In contrast, StabMap performed most efficiently in terms of computation time and memory usage. For cross integration, Concerto and scMDC exhibited the highest runtime while maintaining moderate peak memory usage (Supplementary Fig. 8g,h). MOFA+ also consumed significant computation time and memory, and Multigrate encountered an out of memory error when applied to integrate dataset D56. Again, StabMap was fast and efficient in memory usage in cross integration applications. In spatial registration, most methods run into the out of memory issue due to high memory requirement and SPIRAL was the only method that scale to data with large number cells even though it required long runtime and substantial amount of memory (Supplementary Fig. 8i,j). When a sufficient amount of memory is available, PASTE with centre and pairwise alignment and PASTE2 were computationally efficient to run.

### Impact of clustering algorithms on method evaluation

Given that different clustering algorithms can lead to varying clustering results, we assessed whether the choice of clustering algorithm would influence the ranks of methods in clustering evaluation. Specifically, we compared the overall rankings of methods obtained using three different clustering algorithms of k-means, Leiden, and Louvain, across all integration categories and tasks that included clustering evaluation. The correlations of method rankings among different clustering algorithms are visualised in Supplementary Fig. 9. Our analysis revealed minimal impact on method assessment, as indicated by the high correlations of ranks across different clustering algorithms. For the vertical, diagonal, mosaic, and cross integration categories (Supplementary Fig. 9a-d), most correlations exceeded 0.9. For the feature selection and imputation tasks (Supplementary Fig. 9e,f), the correlations were nearly 1, indicating that different clustering algorithms led to essentially identical assessment results on method performance. These findings demonstrate that our benchmark analyses are robust to the choice of clustering algorithm.

### Method robustness and consistency

To assess the robustness and consistency of the methods, we first performed a stability analysis by randomly excluding approximately 20% of the cell types from the data (repeated 10 times) and examining how each method’s ranking changed relative to the others. This allows us to provide a comparative interpretation of method robustness and consistency that mitigates the lack of interpretation for individual evaluation metrics. Overall, most methods demonstrated robust performance, with relatively small error bars across iterations, indicating consistent rankings (Supplementary Fig. 10). However, a few methods showed notable variability. For instance, in vertical integration of RNA+ADT data modality, Matilda and UINMF exhibited larger error bars compared to other methods, suggesting greater sensitivity to data perturbations (Supplementary Fig. 10a). In feature selection tasks, MOFA+ consistently outperformed other methods with the highest reproducibility, highlighting its robustness across perturbations (Supplementary Fig. 10b). In diagonal integration, scBridge showed stable performance across dimension reduction, clustering, and classification metrics but displayed a larger error bar for batch correction, indicating potential variability in aligning batches when subsets of cell types were excluded (Supplementary Fig. 10c). We also assessed the methods that require random seeds. This analysis complements the evaluation using data perturbation, providing further insights into the sensitivity of methods to stochasticity of algorithms in data integration (Supplementary Fig. 11). We found that variabilities introduced by different random seeds generally are lower than those from data perturbation. Methods such as Matilda that are sensitive to data perturbation show more comparable stability in rankings compared to other methods. These results provide insights into the method robustness and strengthen the comparative analysis of method performance.

## Discussion

We benchmarked 40 methods for vertical, diagonal, mosaic, and cross integration of single-cell multimodal omics data across seven tasks including dimension reduction, clustering, batch correction, classification, feature selection, imputation, and spatial registration. Using a battery of evaluation metrics, we summarise the overall rank score of each method for each integration category and task. This comprehensive ranking provides a detailed assessment of the strengths and weaknesses of methods across distinct evaluation criteria, offering valuable insights into their performance in addressing specific challenges of integration tasks. Overall, the performance of methods is found to be dependent on evaluation metrics, tasks and datasets. These results underscore the importance of evaluating methods using diverse metrics and datasets with different complexity and modality combinations for specific tasks.

In particular, it is important to be aware of the limitation of evaluation metrics. For example, ASW_cellType, iASW, ASW_batch, and PCR cannot be applied to integration methods that generate graph-based outputs. Many evaluation metrics, such as cLISI, rely heavily on accurate cell type annotations which may not always be available. Furthermore, SCS, a spatial registration metric, produces identical assessment scores for methods that rely solely on linear transformations of the data, making it unable to distinguish the relative performance of those methods. These findings highlight the need for careful consideration, selection, and interpretation of evaluation metrics and results to ensure a meaningful method assessment for single-cell multimodal omics data integration.

The benchmark results also revealed a trade-off between preserving biological signals, as reflected by cell type separation, and batch correction. Methods that excel in one aspect may compromise its performance in the other. These findings emphasise the importance of considering specific integration objectives when selecting methods, as the trade-off between different aspects of performance may vary depending on the dataset and analysis priorities. In addition, we found that, for most methods, the performance on cell type classification is primarily influenced by the quality of data integration rather than by the choice of classification model. Indeed, a simple MLP classifier was able to achieve comparable performance to specialised classification models in most cases, suggesting that robust data integration plays a more critical role in downstream classification outcomes.

For the imputation task, we found that methods excelling in clustering and classification metrics do not necessarily perform well in preserving structural features. Likewise, for the feature selection task, approaches that identify cell-type-specific markers often compromise reproducibility when compared to methods that select a single set of markers for all cell types. Consequently, it is essential to consider the specific application, such as prioritising the consistency of feature selection results or the importance of cell-type-specific information, when choosing appropriate methods for these tasks.

Although simulated datasets offer a valuable advantage by providing ground truth that is typically unavailable in real datasets, they may not fully capture the complexity and multidimensional variability inherent in real datasets. Therefore, the performance of integration methods evaluated on the simulated data may not always accurately reflect their performance on real datasets. Indeed, we observed that several methods that performed well on simulated datasets exhibited reduced performance when applied to real datasets. This discrepancy underscores the importance of cautiously interpreting benchmarking results derived from simulations and highlights the necessity of validating method robustness and generalisability using diverse, experimentally derived datasets.

In general, deep learning methods are popular, accounting for 26 out of the 40 integration methods evaluated. Among these, deep learning approaches consistently achieved top performance, securing the highest rank across all data types in both diagonal and cross integration tasks. These methods typically involve multi-layer neural networks, which are computationally intensive and require GPU acceleration to achieve computational efficiency. Additionally, the performance of deep learning methods is often affected by initial values and random seed selection, although the impact of these stochastic factors is generally small.

As a general guide, Fig. 6 summarises the recommendation of top-3 integration methods based on the integration categories and tasks the user aims to accomplish. In particular, the selection of the methods depends on key data characteristics such as whether multiple batches are present, whether all data modalities are present, and if a bridge modality is present to connect datasets. For each category, we recommend three methods based on their overall rank score performance across all datasets for each evaluation criterion, and complemented by considerations of speed and memory usage. To further facilitate an in-depth exploration of method performance on specific tasks, we have also developed a Shiny app (http://shiny.maths.usyd.edu.au/scMultiBench/), which enables users to dynamically and interactively visualise benchmark results. Together, this benchmark study provides a comprehensive overview of computational methods for single-cell multimodal omics data integration, offering valuable guidance to researchers and method developers for applying and developing new methods tailored to specific aspects of data integration. In future work, we aim to address the lack of appropriate and broadly applicable metrics for evaluating integration methods across different tasks.

**Fig. 6:**
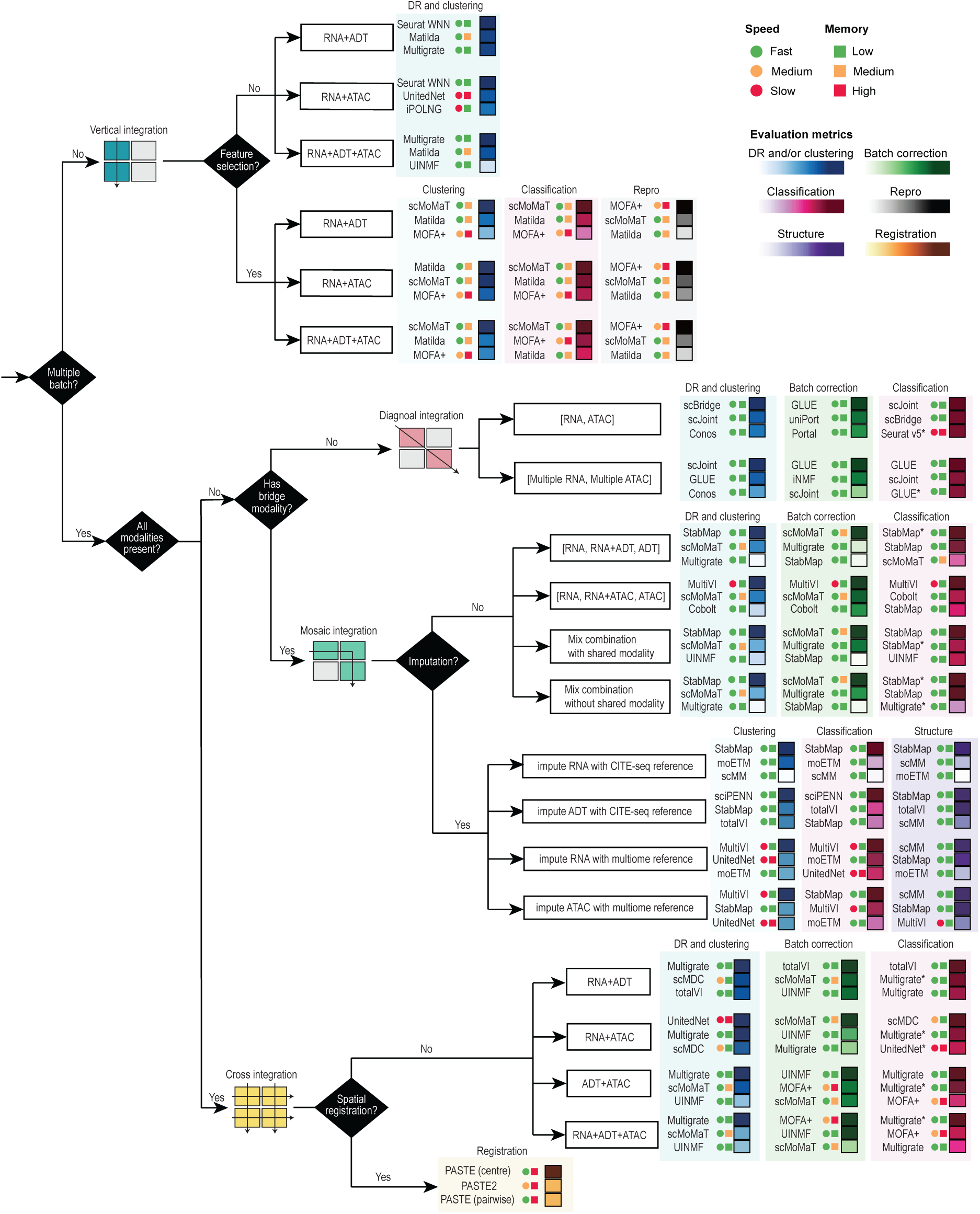
Guidelines for analysing single-cell multimodal omics data. Recommendations of methods were based on data characteristics and the integration categories. The top-3 methods in each specific application and under each category of evaluation metrics were included in the recommendation. The speed and memory usage are colour coded to provide a general guide on method scalability.

## Methods

### Summary of integration methods in each category and task

The evaluation of integration methods from the four integration categories and seven tasks are summarised in Supplementary Fig. 12a-d. Please refer to the Supplementary Notes for the criteria used for selecting integration methods. The details of the 64 real datasets and 22 simulated datasets that are used for the benchmarking are included in the “Datasets and preprocessing” section, and the experimental settings for each of the 40 integration methods are described in the “Single-cell multimodal omics data integration methods” section.

### Datasets and preprocessing

#### Experimental datasets

We create a total of 64 datasets from 21 data sources (see Section “Data availability” in Supplementary Notes) generated using a variety of single-cell multimodal omics technologies. Details of each dataset grouped by integration categories are provided in the Supplementary Notes.

All datasets are listed in Supplementary Table 1, and the datasets employed for each method and task are summarised in Supplementary Fig. 12a-d. The same preprocessing procedure are applied to each dataset including the removal of cell types with fewer than 10 cells, the removal of genes expressed in less than 1% of cells in the RNA data modality, and the filtering of peaks quantified in less than 1% of cells in the ATAC data modality. All features are retained for the ADT data modalities. For multiple ATAC data batches with different genomic ranges, we have followed https://stuartlab.org/signac/articles/merging using ‘reduce’ function from the *GenomicRanges* package and ‘FeatureMatrix’ function from the *Signac* package^54^ to consolidate and quantify the intersecting peaks across all data batches to ensure a cohesive analysis of the data.

#### Data cleaning and reannotation

The reliance on predefined cell types can introduce bias towards methods that resemble those used to produce the original annotations. To minimise the potential bias in the original cell type annotation, we conduct data cleaning and reannotation using AdaSampling^55^ and Annotatability^56^ techniques, follow by manual inspection for additional verification. In AdaSampling, a soft-label learning step is used to train a classification model for inferring mislabelling probability of each cell. Cells with high probabilities of mislabelling are excluded in further iterations of model training. This iterative training process enhances the accuracy of the classification model by systematically improving the quality of the cell type labels used for training. For each cell, the final trained model is used to either confirm its original cell type annotation or identify potential mislabelling and possible correction for the cell. Similarly, Annotatability refines cell type annotation with a deep neural network (DNN) by monitoring the confidence (quantified as the average probability of annotations) and the variability (quantified as the standard deviation for the probability of annotations) over training epochs. Cells are then categorised as correctly annotated, erroneously annotated, or ambiguous based on their confidence and variability. Possible corrections for cells in the erroneously annotated category are given by DNN trained solely on cells from the correctly annotated category.

The data cleaning and reannotation procedure is summarised in Supplementary Fig. 12e. Specifically, we reannotate cells to their newly nominated cell types if both AdaSampling and Annotatability identified them as erroneously annotated and nominated them to the same new cell type. Cells that are identified as erroneously annotated by both methods, but the newly annotated cell type disagree for the two methods are removed. Finally, cells that are deemed as corrected annotated if both methods classify them to their original cell type. The final dataset after the cleaning and reannotation procedure is manually validated by inspecting the expression of differentially expressed genes from each cell type and the annotated cells to verify if they are at a comparable level.

#### Simulation datasets

While the application of the above reannotation procedures may help reduce the potential cell annotation bias from the predefined cell types of each data source, simulation datasets can also be used as an alternative for method evaluation. We use a generalised version of SymSim^57^ to simulate single-cell multimodal omics for method evaluation. Specifically, we simulate multiome data with RNA and ATAC modalities by following the tutorial (https://github.com/PeterZZQ/Symsim2) provided by the scDART^58^ using the functions ‘SimulateTrueCounts’, ‘True2ObservedCounts’, and ‘DivideBatches’ in the package. For the simulation of ADT data modality, we follow the instruction outlined in scMoMaT^21^. This involves choosing a reference ADT count dataset to establish a count distribution, from which we then simulate ADT counts that exhibit a positive correlation with gene expression data. For integration methods that require gene activity scores as input, we simulate the gene activity score using the same procedure as in simulating the ADT modality. After simulating all paired modalities, we create various modality combinations according to the integration categories. We create a total of 22 simulated datasets. Details of each dataset grouped by integration categories are provided in the Supplementary Notes.

### Single-cell multimodal omics data integration methods

In this section, we outline the single-cell multimodal omics data integration techniques included in this benchmark study. Count data is typically utilised as the input, unless otherwise indicated. For diagonal integration, we select methods specifically designed for aligning RNA and ATAC data modalities. Some methods require using original ATAC peaks directly as input, while others require peak data converted into gene activity scores. To facilitate this, we implement a cohesive pipeline, using the ‘GeneActivity’ function from the Signac package (v1.14.0), to transform the ATAC matrix from peak level quantifications to gene activity scores. Note that we use the ‘CreateGeneActivityMatrix’ function in the Seurat package to convert peak level quantifications into gene activity scores for the ‘P0_BrainCortex’ sample of SNARE-seq data as fragment files are not available. For diagonal integration, we use the gene activity scores from ATAC data as default unless indicated otherwise. For vertical, mosaic, and cross integration, we use the peak data of ATAC as default unless indicated otherwise. The appropriate data combinations and formats for each method are included in Supplementary Table 2. Details for each method are provided in the Supplementary Notes.

### Evaluation metrics

Here we summarise the metrics used for evaluating methods on single-cell multimodal omics data integration. The definition of each metric is provided in the **Supplementary Notes**. For each metric, the advantages, disadvantages, requirements for cell type and batch labels, applicability to graph or embedding outputs, and other information are included in Supplementary Table 3.

#### Dimension reduction

Biological conservation via Graph cell-type local inverse Simpson’s index (cLISI) score, and dimension reduction scalability via computational time (CT) and memory usage (MU).

#### Batch correction

Batch correction quality via k-nearest neighbour batch effect test (kBET) score; graph integration LISI score (iLISI); adjusted Rand index by batch (ARI_batch); normalised mutual information by batch (NMI_batch); average Silhouette width by batch (ASW_batch); graph connectivity (GC); and principal component regression (PCR) score.

#### Clustering

Clustering quality evaluation via adjusted Rand index by cell type (ARI_cellType); normalised mutual information by cell type (NMI_cellType); average silhouette width by cellType (ASW_cellType); isolated label score of F1 (iF1); and isolated label score ASW (iASW).

#### Classification

Classification performance evaluated via the overall classification accuracy (OCA); average classification accuracy (ACA); specificity (Spec) and sensitivity (Sens); and F1-score (F1).

#### Feature selection

Feature specificity and reproducibility via marker overlap among different cell types (MO) and marker correlation among downsampled data (MC); and downstream analysis metrics including clustering and classification metrics applied to top features.

#### Imputation

Imputation data structure metrics including standardized MSE imputation error with ground truth data (sMSE); preservation of feature correlation structure (pFCS); and preservation of differential expression statistics (pDES). Imputation downstream analysis metrics including clustering and classification metrics applied to imputed data.

#### Spatial registration

Spatial registration quality metrics include Label Transfer Adjusted Rand Index (LTARI); Pairwise Alignment Accuracy (PAA); and Spatial Coherence Score (SCS). Spatial registration scalability metrics include computational time (CT_SR) and memory usage on spatial registration (MU_SR).

### Evaluation pipelines

The following sections describe the seven task evaluation pipelines. The details of the datasets, integration methods, tasks and evaluation metrics involved are described in the corresponding sections above. The code for each pipeline is provided at https://github.com/PYangLab/scMultiBench/tree/main/evaluation_pipelines. The detailed description of the algorithms and pseudocode are included in Supplementary Notes.

#### Dimension reduction evaluation pipeline

The dimension reduction task evaluation pipeline is summarised in Supplementary Fig. 13a and by Algorithm 1 (Supplementary Notes). The input data can be either single-batch multimodal count data or multi-batch multimodal count data, and the multimodal integration methods typically generate low-dimensional embeddings/graphs that capture the multimodal characteristics of individual cells. The quality of the embeddings/graphs generated by the integration methods can be evaluated using the cLISI score. We also assess the scalability of these integration methods by analysing the computational time (CT) and memory usage (MU). Specifically, with our dimension reduction evaluation pipeline, we analyse 18 vertical integration methods on 23 real datasets and 6 simulated datasets, 14 diagonal integration methods on 14 real datasets and 4 simulated datasets, 7 mosaic integration methods on 13 real datasets and 4 simulated datasets and 11 cross integration methods on 9 real datasets and 8 simulated datasets (Supplementary Fig. 13a).

#### Batch correction evaluation pipeline

Batch correction is carried out using various multi-batch integration methods, including 14 diagonal integration methods on 14 real datasets and 4 simulated datasets, 7 mosaic integration methods on 13 real datasets and 4 simulated datasets and 11 cross integration methods on 9 real datasets and 8 simulated datasets. These methods can generate batch-corrected low-dimensional embeddings/graphs, which is evaluated using the 7 batch correction metrics. Note that PCR is calculated only for cross integration because it requires unintegrated data. Cross integration is the only category where there is unintegrated data for a specific modality across all batches. Supplementary Fig. 13b illustrates the evaluation pipeline for the batch correction task.

#### Clustering evaluation pipeline

The clustering evaluation pipeline is summarised in Supplementary Fig. 13c. In particular, clustering is performed on the low-dimensional integrated embeddings/graphs generated from the dimension reduction methods using Leiden clustering. All integration methods and datasets from the dimension reduction evaluation pipeline is assessed, including 18 vertical integration methods on 23 real datasets and 6 simulated datasets, 14 diagonal integration methods on 14 real datasets and 4 simulated datasets, 7 mosaic integration methods on 13 real datasets and 4 simulated datasets and 11 cross integration methods on 9 real datasets and 8 simulated datasets. The quality of the clustering results is measured using the 5 clustering evaluation metrics, including ARI_cellType, NMI_cellType, ASW_cellType, iF1 and iASW.

#### Classification evaluation pipeline

**Supplementary** Fig. 13d summarises the workflow of the classification evaluation pipeline. Similar to the batch correction pipeline above, classification is also performed and evaluated on the integrated low-dimensional embeddings generated from the dimension reduction methods, with the focus on the multi-batch integration methods, including the 14 diagonal integration methods on 14 real datasets and 4 simulated datasets, the 7 mosaic integration methods on 13 real datasets and 4 simulated datasets and the 11 cross integration methods on 9 real datasets and 8 simulated datasets. All datasets included in this task contain multiple batches and for each dataset and a specific method, each data batch is used as a reference, with the remaining batches as queries to assess prediction results using a simple multi-layer perceptron (MLP) classifier. If a method also provides its own classification model, we also include the classification results using its own model. Finally, we summarise the results from across all classification combinations for each dataset and method.

#### Feature selection evaluation pipeline

Summarised in the evaluation pipeline in Supplementary Fig. 13e, we focus on the 3 integration methods that enable feature selection, including Matilda, scMoMaT and MOFA+. The quality of feature selection is evaluated on the 23 real datasets and 6 simulated datasets and using the aforementioned 5 classification metrics, 5 clustering metrics, and the feature specificity and reproducibility metrics (i.e. MO and MC). In particular, we train a simple MLP classifier and conduct 5-fold cross-validation for evaluating the utility of selected features for classifying cell types and also apply the Leiden clustering pipeline as in the clustering task to evaluate their utility in cell clustering.

#### Imputation evaluation pipeline

Supplementary Fig. 13f illustrates the imputation evaluation pipeline. We focus on the 7 mosaic integration methods applicable to imputation. This evaluation involves using reference and query data generated from datasets D51-57. In particular, we randomly select one batch as the reference data and then select another batch as query but only keep one of data modality in the query dataset. This allows the missing data modality in the query data to be used as the ground truth for evaluating the quality of the imputed data. During the imputation, we utilise the reference data to impute the missing data modalities in the query data and then compare the imputed data with the ground truth data. The quality of the imputed data is quantified using 3 imputation data structure metrics (i.e. sMSE, pFCS, pDES) as well as downstream analysis metric, including 5 classification metrics and 5 clustering metrics. Specifically, for the classification assessment, we first train a simple MLP classifier using the reference data subset by the features that are included in the imputed data (denoted by * in Supplementary Fig. 13f) and then classify the cells using the imputed data. This approach allows for the assessment of imputed data using the 5 classification metrics.

#### Spatial registration evaluation pipeline

The spatial registration evaluation pipeline focuses on the 4 integration methods applicable to this task using the 5 spatial datasets with multiple slices, with each spatial slice consisting of the gene expression and spatial coordinates. Supplementary Fig. 13g illustrates the evaluation schematic which uses multiple spatial slices as the input. After conducting spatial registration via the integration methods, these slices are aligned into a consistent coordinate system (i.e. aligned coordinates). For evaluation, we employ 3 established metrics from existing literature, including LTARI, PAA, and SCS. For LTARI, we perform Leiden clustering on all slices independently to obtain cluster labels for each slice. We next choose one slice as the reference, while the remaining as queries and transfer labels from the reference to the queries using 1-nearest neighbours based on the aligned coordinates. Finally, we apply the ARI metric to the transferred label and ground truth to calculate LTARI. For PAA and SCS, they are calculated based on the reference labels and the labels transferred to the query data.

Besides, we also apply 2 spatial registration scalability metrics for the evaluation, including CT_SR and MU_SR. As described in the evaluation metrics section, we randomly sample from dataset 61 to create 11 sub-datasets with different cell numbers ranging from 500 to 100,000, based on which the runtime (CT_SR) and memory usage (MU_SR) are evaluated.

### Other benchmark considerations

#### Robustness and consistency assessment

We randomly remove a subset of cell types (∼20%) from datasets to assess the robustness of methods on data perturbation. The random removal of cell types is repeated 10 times and for each iteration, we evaluate the performance of the integration using metrics for their respective tasks. The change of performance on perturbed datasets is used to assess the robustness of each method on all integration categories and tasks. We also assess the impact of methods that rely on random initialisation by randomising initial values/seeds and repeating these methods (10 times) to evaluate their integration performance using metrics for their respective tasks. Finally, we assess the consistency of the methods by grouping and ranking their performance results by technologies and/or platforms. This allows us to quantify the change of their performance across different technologies and platforms.

#### Clustering algorithms and parameters

We use popular clustering algorithms of k-means, Leiden, and Louvain, for tasks that require clustering to evaluate method performance. Specifically, for k-means, we set k, the number of clusters, to the actual number of cell types in each dataset. For Leiden and Louvain clustering, we employ the ‘get_N_clusters’ function from the Sinfonia package^59^ to automatically select the resolution parameters that align the number of clusters to the number of cell types in each dataset. This allows us to quantify the performance of integration methods across different clustering algorithms.

### R Shiny application for interactive exploration of benchmark results

We create an R Shiny application to host the benchmark results for interactive exploration, enabling the users to select specific integration categories, tasks, evaluation metrics, and datasets of interest and dynamically display the relative performance of methods across the selections. The application will also enable the inclusion of new methods and datasets as they become available, ensuring up-to-date communication of benchmark results.

### Guideline and Recommendations

We provide a decision-tree-style guideline for each integration and task category as the final recommendation for method selection. Specifically, the decision-tree-style guideline includes scenario-specific recommendations for methods, taking into consideration data characteristics and computational efficiency. Each branch of the decision tree displays three recommended methods tailored to the specific integration category and task. These methods are accompanied by key performance metrics benchmarked in this study represented using colour boxes. The structured guidelines facilitate clear and comprehensive method recommendations for users.

## Data availability

All real single-cell and spatial transcriptomics datasets used in this benchmark were obtained from publicly available repositories, as described in Supplementary Table 4. These include CITE-seq datasets from ArrayExpress (E-MTAB-10026) and GEO (GSE166489, GSE164378, GSE194122), Zenodo (https://zenodo.org/records/6348128), and Software Heritage (https://archive.softwareheritage.org/browse/revision/1c7fcabb18a1971dc4d6e29bc3ed4f6f36 b2361f/); 10x multiome datasets from GEO (GSE194122, GSE205117, GSE204684) and Zenodo (https://zenodo.org/records/6348128); SHARE-seq and SNARE-seq datasets from GEO (GSE140203, GSE126074); ASAP-seq, DOGMA-seq, and TEA-seq datasets from GEO (GSE156478, GSE158013); and spatial transcriptomics datasets including Visium (https://zenodo.org/records/6334774, https://github.com/raphael-group/paste_reproducibility/tree/main/data/DLPFC), Xenium (https://www.10xgenomics.com/products/xenium-in-situ/preview-dataset-human-breast), Stereo-seq (https://db.cngb.org/stomics/flysta3d/), MERFISH (https://cellxgene.cziscience.com/collections/31937775-0602-4e52-a799-b6acdd2bac2e), and Spatial ATAC-RNA (GEO: GSE205055). The processed input datasets used for benchmarking are available in a publicly accessible Figshare repository (https://figshare.com/articles/dataset/datasets/29035586?file=54438737).

## Code availability

We have uploaded the source code to a GitHub repository at https://github.com/PYangLab/scMultiBench, including scripts for running the benchmarking methods, the evaluation pipeline, and generating the figures associated with the paper. The code is also archived and available via Zenodo at https://doi.org/10.5281/zenodo.15385334.

## Acknowledgements

The authors thank all their colleagues, particularly at the School of Mathematics and Statistics, The University of Sydney, and Sydney Precision Data Science Centre for their support and intellectual engagement. This work was supported by a National Health and Medical Research Council (NHMRC) Investigator Grant (1173469) to P.Y.. S.G. was supported by an Australian Research Council DECRA Fellowship (DE220100964).

## Author contributions statement

CL, HJK, and PY conceived the study. PY supervised the study and CL, SD, SL, and DX performed the initial feasibility analysis of the study. CL led the benchmark with major contributions from SD and PY, and additional contributions from HJK, SL, DX and SG. CL, SD, and PY drafted the manuscript with input from all authors. All authors read, edited, and approved this work.

## Competing interests statement

The authors declare they have no competing interests.

## Notes

### Competing Interest Statement

The authors have declared no competing interest.

### Summary of Updates

To include experimental results from the benchmark study.

